# CIPK-B is essential for salt stress signalling in *Marchantia polymorpha*

**DOI:** 10.1101/2022.08.22.504506

**Authors:** Connor Tansley, James Houghton, Althea M. E. Rose, Bartosz Witek, Rocky D. Payet, Taoyang Wu, J. Benjamin Miller

## Abstract

- Calcium signalling is central to many plant processes, with families of calcium decoder proteins having expanded across the green lineage and redundancy existing between decoders. The liverwort *Marchantia polymorpha* has fast become a new model plant, but it is unclear what calcium decoders exist in this species.
- We have performed phylogenetic analyses to identify the Calcineurin B-Like (CBL) and CBL-Interacting Protein Kinase (CIPK) network of *M. polymorpha*. We analysed CBL-CIPK expression during salt stress, and determined protein-protein interactions using yeast two-hybrid and bimolecular fluorescence complementation. We also created genetic knockouts using CRISPR/Cas9.
- We confirm that *M. polymorpha* has two CIPKs and three CBLs. Both CIPKs and only one CBL show salt-responsive transcriptional changes. All *M. polymorpha* CBL-CIPKs interact with each other *in planta*. Knocking out *CIPK-B* causes increased sensitivity to salt suggesting that this CIPK is involved in salt signalling.
- We have identified CBL-CIPKs that form part of a salt tolerance pathway in *M. polymorpha*. Phylogeny and interaction studies imply that these CBL-CIPKs form an evolutionarily conserved Salt Overly Sensitive (SOS) pathway. Hence, salt responses may be some of the early functions of CBL-CIPK networks and increased abiotic stress tolerance required for land plant emergence.

## INTRODUCTION

Salinization of arable land is a leading threat to sustainable global food security and plants demonstrate reduced yield at concentrations as low as 40 mM NaCl (Munns, 2005). It is estimated that 33% of irrigated land globally is afflicted by high salinity, and, with salinized areas increasing by 10% a year, salinization is predicted to affect more than 50% of arable land by 2050 (Jamil *et al*., 2011). To prepare for increasing agricultural land salinization, knowledge of abiotic stress response systems in plants has to be better understood. Release or influx of calcium at different subcellular and tissue localisations acts as an essential secondary messenger in plants, where it is important for cell polarity, directional growth, fertilisation and responses to abiotic and biotic stresses (Knight & Knight, 2001; Li *et al*., 2006; Boudsocq *et al*., 2010; Boudsocq & Sheen, 2013; Edel & Kudla, 2015). Hence, understanding calcium signalling responses under salt stimuli is essential to developing new approaches to maintain or increase current global food supply in response to increasing pressures from abiotic stress.

Salt stress initiates calcium signals in plants (Knight *et al*., 1997; Choi *et al*., 2014; Huang *et al*., 2017) that are decoded by various calcium-binding proteins (Allan *et al*., 2022). These decoder proteins detect calcium fluctuations by binding calcium through intrinsic helix-loop-helix domains, known as EF-hands, and fall broadly into sensor-relay and sensor-responder types. Sensor-responders, such as Calcium-Dependent Protein Kinases (CDPKs), detect calcium changes through EF-hands and then directly phosphorylate downstream proteins via intrinsic kinase domains (Bredow & Monaghan, 2022). Sensor-relays, such as Calcineurin B-Like proteins (CBLs) and Calmodulins (CaMs), have no intrinsic kinase activity but undergo a conformational change upon binding of calcium which allows them to interact with downstream partners to regulate the activity of other signalling proteins (Zhang & Lu, 2003; Kolukisaoglu *et al*., 2004; Hashimoto *et al*., 2012). CBL-Interacting Protein Kinases (CIPKs) are essential to transduce calcium signals in plants through the interactions that they form with CBLs (Albrecht *et al*., 2001).

CBLs and CIPKs are important for salt stress signalling and were initially characterised in the Salt Overly Sensitive (SOS) pathway of *Arabidopsis thaliana*. In this pathway, SOS3 (AtCBL4) is activated by a salt-induced calcium signal in roots. SOS3 then activates SOS2 (AtCIPK24), which transduces the signal by phosphorylating the Na^+^/H^+^ antiporter SOS1 (Qiu *et al*., 2002; Ji *et al*., 2013) to increase Na^+^ extrusion, thereby stopping Na^+^ from entering the transpiration stream (Quan *et al*., 2007; Quintero *et al*., 2011). The interaction between SOS3 and SOS2 is dependent on the SOS2 NAF domain and removal of this domain renders SOS2 auto-active but less capable of activating SOS1 (Albrecht *et al*., 2001; Quintero *et al*., 2011). The SOS pathway is in fact more complex as ScaBP8 (AtCBL10) functions in place of SOS3 (AtCBL4) in the shoot (Quan *et al*., 2007) to interact with SOS2 and activate SOS1 for salt extrusion, and AtCIPK8 can functionally replace SOS2 in the same pathway (Yin *et al*., 2020). This discovery of many CBL-CIPKs that can effectively fulfil the same role demonstrates the level of functional redundancy between these proteins and highlights the complexity of calcium signalling during salt stress.

Functional redundancy is common in calcium decoders and can arise from whole genome duplication, but maintenance of selection is required for redundancy not to be lost. Analysis of the evolution of calcium signalling components has demonstrated that the increase of calcium encoding machineries (such as ion channels) in early land plants was followed by a subsequent increase in decoder components (Edel *et al*., 2017). To increase the number of calcium decoders, mechanisms to increase gene number would have been required during evolution. Indeed, whole genome duplication and hybridisation are common in later parts of the green lineage (Clark & Donoghue, 2018). Thus far, studies have focussed on these later diverging groups, such as *A. thaliana* and crops, which have functional redundancy likely due to these whole genome duplication events and maintenance of selection. Studies have analysed the CBL-CIPK network of the moss *Physcomitrium patens* (*Physcomitrella*) (Kleist *et al*., 2014), but even *P. patens* has a rather extensive network of six CBLs and eight CIPKs. One of these CBLs (PpCBL6) is not expressed, and an additional CBL (PpCBL5) and CIPK (PpCIPK8) are likely pseudogenes. Together this implies that there are only four PpCBLs and seven PpCIPKs that are functional and demonstrates the need for maintaining selection to keep redundancy. Of these remaining *P. patens* CBLs and CIPKs, most form cognate pairs at genomic loci, which suggests that CBLs and CIPKs duplicated in the whole genome duplication events proposed in *P. patens* (Rensing *et al*., 2007; Kleist *et al*., 2014). Most of the PpCIPKs fall into the “algal-type” clade, which includes AtCIPK24 (SOS2) and AtCIPK8, and hence it could be proposed that these CIPKs function in salt tolerance responses. Indeed, PpCIPK1 has been characterised to function in salt tolerance (Xiao *et al*., 2021). However, bryophytes diverged from the rest of the green lineage >400 million years ago and it is therefore essential to characterise multiple bryophytes before salt stress tolerance can be proposed as an ancient function of CBL-CIPK networks.

*Marchantia polymorpha* has been recently established as another bryophyte model organism and a range of molecular biology and genetic tools have been developed for this species, including *Agrobacterium*-mediated transformation and CRISPR/Cas9 mutagenesis (Bowman *et al*., 2017; Sauret-Güeto *et al*., 2020). Key plant signalling mechanisms are conserved in *M. polymorpha*, including both auxin and cytokinin signalling albeit with fewer genes involved (Flores-Sandoval *et al*., 2015; Kato *et al*., 2015; Aki *et al*., 2019). Similarly, jasmonate signalling components have been discovered in *M. polymorpha* (Monte *et al*., 2019). Whilst *M. polymorpha* can respond to ethylene, it cannot synthesise ethylene and instead produces the precursor 1-aminocyclopropane-1-carboxylic acid (ACC) (Li *et al*., 2020). Although many of the key plant hormone signalling components are present and functional in *M. polymorpha*, calcium signalling components have not yet been explored. Furthermore, the similarities and differences between signalling in *M. polymorpha* and other land plants highlight the need to understand signalling, including calcium signalling, in more than one bryophyte. This is particularly necessary to also propose plausible functions for calcium signalling in the last universal common ancestor of land plants. Here, we identify the CBL-CIPK network of calcium decoders in *M. polymorpha* and show that specific CBL-CIPK protein-protein interactions are not found in this bryophyte species. We also demonstrate that *M. polymorpha* has a CIPK (MpCIPK-B) with a salt stress signalling role, specifically in response to ionic stress rather than osmotic stress. We propose that *M. polymorpha* has an evolutionarily conserved SOS pathway and that salt responses may be some of the early functions of CBL-CIPK networks in land plants.

## MATERIALS AND METHODS

### Phylogeny

*Arabidopsis thaliana* CBL and CIPK protein sequences were used as a BLAST query against predicted protein sequences from *Marchantia polymorpha* (version 5.0, marchantia.info) with an e-value cut off of 1e-50. Sequences were filtered for unique IDs and full length sequences were retrieved. *Physcomitrium patens, Klebsormidium nitens, A. thaliana* and *M. polymorpha* sequences (Table S1) were used for multiple alignment with MAFFT (version 5.0) followed by trimming with trimAl (version 1.3) and maximum likelihood trees were generated with IQ-Tree (1.6.12) with 1000x bootstrapping (model: JTTDCMut+G4). Trees were visualised with the Interactive Tree of Life (iTOL) (Letunic & Bork, 2021).

### Plant growth and phenotyping

Four accessions of *M. polymorpha* were used in this study: Takaragaike-1 (male), Takaragaike-2 (female), Cambridge-1 (male), Cambridge-2 (female), supplied by the Haseloff laboratory (University of Cambridge). *M. polymorpha* was maintained on 0.5 MS media with 1% sucrose and 0.8% agar (pH 6.0) and grown under a 16 hour photoperiod at 23 °C. Phenotyping was carried out with adult plants (>3 weeks) by taking ~5×5 mm cuttings and placing them on MS plates supplemented with the indicated concentrations of NaCl or sorbitol. Five cuttings were used for each replicate and grown for 7 days. A pooled fresh weight was taken for the five plants and the plant material was then snap frozen in liquid nitrogen for subsequent RT-qPCR analysis or used fresh to assess chlorophyll content.

### Expression analysis by RT-qPCR

RNA was extracted from plant material using Qiagen RNeasy Plant Minikit following the manufacturer’s instructions. DNA was removed by DNase treatment (AMbion Turbo DNase). RNA quality was confirmed by measuring the absorbance at 260 and 280 nm using a NanoDrop™ 8000 spectrophotometer (NanoDrop Technologies). To test for the absence of contaminating genomic DNA, a PCR reaction was performed using the GoTaq^®^ G2 Green Master mix (Promega) and primers specific for the housekeeping gene *Actin1 (ACT1;* Table S2). The PCR conditions were as follows: 30 s at 98 °C, followed by 30 cycles of 10 s at 98 °C, 20 s at 53 °C and 45 s at 72 °C, followed by 5 min at 72 °C. Genomic DNA from wildtype plants was used as a positive control. The absence of contaminating genomic DNA was confirmed by the lack of a 154 bp band when the PCR product was analysed by agarose gel electrophoresis. Approximately 1 μg of purified RNA from each sample was also analysed by agarose gel electrophoresis to confirm the presence of the 18S and 28S rRNA double bands. Complementary DNA was synthesised using Superscript™ II (Invitrogen) and oligo-dT17 with at least 500 ng of RNA as input. RNase activity was inhibited during cDNA synthesis using 1 μl RNAsin. Quantitative RT-PCR was carried out using SYBR Green (Jumpstart Taq ReadyMix: Sigma-Aldrich) in 10 μl volumes for 3 biological and 3 technical replicates with 1:10 diluted cDNA. Measurements were taken on AriaMX Real-Time qPCR system (Agilent).

*Actin1* and *Adenine phosphoribosyl transferase (APT)* were used as housekeeping genes. Conditions for RT-qPCR were as follows: 95 °C for 4 minutes; 40 cycles of 94 °C for 30 seconds, 55 °C for 30 seconds, 72 °C for 30 seconds; followed by cooling and melt curve. Results for each RT-qPCR reaction were expressed as threshold cycle (Ct) values. Sequences of the gene-specific primers are listed in Table S2. Efficiencies of primer pairs were tested using a dilution series of 10^-1^-10^-5^ ng/μl of each gene-specific PCR product and confirmed as 90-110%. It was also confirmed that primers did not amplify control samples without cDNA template. Ct values for each gene were averaged across the technical replicates for each cDNA sample. Ct values for the two housekeeping genes (*ACT1* and *APT*) were averaged and used as a control. The fold induction (relative expression) for each biological replicate was calculated for treated samples relative to untreated samples using the Pfaffl method (Pfaffl, 2001) and primer efficiencies calculated for each RT-qPCR plate (as described above). The final fold induction was calculated by averaging results from all three biological replicates.

### Golden Gate cloning

All constructs were assembled using the MoClo Golden Gate cloning system (Weber *et al*., 2011) according to Feike *et al*. (2019). Briefly, Level 0 modules were ordered as synthesised DNA parts (Invitrogen GeneArt) or domesticated from cDNA clones to remove internal BsaI and BpiI restriction enzyme sites using overlap extension PCR or PCR amplification and scarless Golden Gate assembly. Level 1 parts were assembled in one-step restriction-ligation reactions using BsaI, transformed into chemically competent *E. coli* DH5α cells and selected with 100 μg/ml ampicillin and blue/white screening. Level 2 multigene constructs were made using BpiI in one-step restriction-ligation reactions, transformed into chemically competent *E. coli* DH10B cells and selected with 25 μg/ml kanamycin and red/white screening. All final Golden Gate constructs (Table S3) were validated by colony PCR using gene-specific primers, diagnostic restriction enzyme digestions and Sanger sequencing. Constructs for bimolecular fluorescence complementation (BiFC), split luciferase assays and CRISPR/Cas9 mutagenesis were transformed into *Agrobacterium tumefaciens* strain GV3101, with resulting colonies selected by colony PCR using gene-specific primers.

### Yeast two-hybrid assays

*Saccharomyces cerevisiae* was transformed following the lithium acetate transformation procedure (Gietz & Woods, 2002). Yeast strain AH109 was transformed with constructs containing the GAL4-BD fused to the N-terminus of each CBL (based on pDEST-GBKT7), and strain Y187 was transformed with constructs containing the GAL4-AD fused to the N-terminus of each CIPK (based on pDEST-GADT7). AH109 and Y187 strains were mated to generate diploid yeast strains containing different pairs of GAL4-BD-CBL and GAL4-AD-CIPK constructs. Five biological replicates from each mating were taken and yeast colony PCR, with gene-specific primers, was used to verify the presence of both constructs. To determine protein-protein interactions, yeast growth was assessed on synthetic dropout (SD) media lacking Leu and Trp (SD-LW; non-selective growth on control plate) and SD media lacking Leu, Trp and His (SD-LWH; selective growth for interaction test). Diploid yeast cells were grown in YPAD at 30 °C overnight, then diluted to an OD_600_ of 1.5, and a 1:10 serial dilution was spotted onto control and interaction test plates. Yeast growth was assessed after 4 days growth at 30 °C. Three technical replicates were performed, with all replicates showing similar results.

### Western blotting

The five independent biological replicates used for each yeast two-hybrid interaction test were grown at 30 °C in a 50 ml selective culture (SD-LW with 100 μg/ml carbenicillin) until an OD_600_ of 0.4-0.6 was reached. The cells were pelleted by centrifugation (10 minutes, 1000 rpm), washed by re-suspension in 20 ml of water and re-pelleted again (10 minutes, 1000 rpm). OD units were calculated by multiplying the final OD_600_ by the ml of culture. Per OD unit, 10 μl of Laemelli buffer was added to the pelleted cells to make a normalised re-suspension for each interaction. The yeast cells and Laemelli buffer were then boiled at 95 °C for 30 minutes to lyse the cells. SDS-PAGE was performed on the samples using a 10% resolving gel run at 130 V for 2 hours. The proteins were transferred to PVDF membrane via wet transfer at 100 V for 80 minutes at 4 °C. Membranes were blocked with TBS-T containing 5% milk for 1 hour and then washed with TBS-T. Membranes were then incubated overnight at 4 °C with 10 ml TBS-T containing either anti-c-myc-peroxidase antibody (1:10,000; Sigma, A5598), or anti-HA primary antibody (1:10,000; Sigma, H6908). Anti-HA gels were washed 3 times with 20 ml TBS-T and then incubated with peroxidase-conjugated secondary antibody (1:10,000; Sigma A0545) in TBS-T for 1 hour at room temperature. Peroxidase activity was detected using ECL Select Western Blotting Detection Reagent (Cytiva, Amersham) following the manufacturer’s instructions.

### Protein-protein interaction tests in *Nicotiana benthamiana*

*Nicotiana benthamiana* plants were grown at 23 °C under long-day conditions in compost for 4-5 weeks prior to infiltration. Plant transformation was performed using *Agrobacterium tumefaciens* strain GV3101 transformed with constructs of interest via freeze-thaw transformation or electroporation. Bimolecular fluorescent complementation constructs contained the C-terminal half of Venus fused to each CIPK (Venus^C^-CIPK), or the N-terminal half of Venus fused to each CBL (CBL-Venus^N^). Co-infiltration with two *A. tumefaciens* strains (GV3101), diluted to OD_600_ of 0.5, was performed to assess CBL-CIPK interactions in a pairwise manner. Split luciferase complementation assay constructs contained the SmBiT NanoLuc luciferase (Promega) fused to the CIPK of interest (SmBiT-CIPK), the LgBiT NanoLuc luciferase (Promega) fused to the CBL of interest (CBL-LgBiT), and a GUS transformation marker. Protein-protein interactions were assessed 2-3 days post infiltration with *A. tumefaciens*. Bimolecular fluorescence complementation was assessed using a Zeiss Leica Airyscan LSM880 with 488 nm excitation, and emission detected from 490-543 nm. Images were analysed with ZEN Black 2.1.fluorescence microscopy software as described above. For split luciferase assays, luciferase activity measurements were performed as described by Feike *et al*. (2019), although using Nano-Glo^®^ Live Cell Assay System kit (Promega) containing substrate, dilution buffer and water in a 1:50:49 ratio and a Hidex Sense microplate reader.

### CRISPR/Cas9 mutagenesis

The CRISPR/Cas9 construct contained the Cas9 enzyme (NLS-pcoCas9), two single guide RNAs designed to target the first exon of *CIPK-B* (expressed from the *M. polymorpha* U6 promoter), and a hygromycin resistance cassette (p35S-hptI-tNOS) for selection of positive transformants. *M. polymorpha* transformation was carried out using *A. tumefaciens* strain GV3101. Cam-2 gemmae were co-cultivated with *A. tumefaciens* for 3 days in infiltration buffer (10 μM MgCl_2_, 10 μM MES buffer, 200 μM acetosyringone) with shaking (200 rpm) in the dark at 23 °C. Positive transformants were selected on hygromycin (10 μg/ml), grown to the T1 generation by propagation of gemmae, and confirmed to contain a deletion in *CIPK-B* by PCR amplification of exon 1 from extracted genomic DNA. Sanger sequencing was performed on PCR products to determine the precise mutation present in each *cipk-b* mutant line.

### Chlorophyll content

Chlorophyll content was assessed using an adapted protocol from Caesar *et al*. (2018). Briefly, 10 μl of 200 mM Na_2_CO_3_ was added to weighed fresh plant tissue, followed by 500 μl DMSO. The plant tissue was then homogenised using a micro-pestle and incubated at 65 °C for 90 minutes. Another 500 μl DMSO was added and the incubation repeated. The samples were then centrifuged (5000 rpm for 10 minutes) and 200 μl of each sample was added to a clear 96 well plate in triplicate. The absorbance was then measured at 648, 665, and 700 nm using a Hidex Sense microplate reader. In the case of the 665 nm absorbance reading being greater than 0.8, the sample was diluted in a 1:1 ratio of DMSO to sample. Chlorophyll content was calculated following the equation:

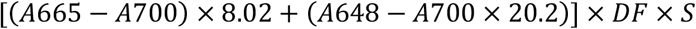

where DF is the dilution factor (usually 1) and S is the amount of solvent (in this case 0.2 ml), and then normalised per unit fresh weight.

## RESULTS

### *Marchantia polymorpha* has a simplified CBL-CIPK network

To determine the complexity of the CBL-CIPK network in *Marchantia polymorpha*, we performed phylogenetic analyses to identify CBL-CIPK proteins in the version 5.0 release of the *M. polymorpha* genome. Our analysis identified three CBLs in *M. polymorpha* with representation in the two main types of CBLs (Fig. 1a). MpCBL-A was grouped with *A. thaliana* CBLs known to localise to the tonoplast, but did not have the motif proposed to confer tonoplast localisation (Batistič *et al*., 2012; Tang *et al*., 2012). MpCBL-A also did not contain the MGCxxS/T myristoylation and palmitoylation motif for localisation to the plasma membrane. MpCBL-B/C both fall into the group of AtCBLs known to localise to the plasma membrane and have the required MGCxxS/T motif for localisation. The three MpCBLs all contain the 14 amino acid-long first EF-hand, which is a key feature of CBLs, as well as the FISL motif which is known to be phosphorylated in CIPK interactions and therefore it is likely that this regulation via phosphorylation is present in early-diverging land plants.

**Figure 1:**
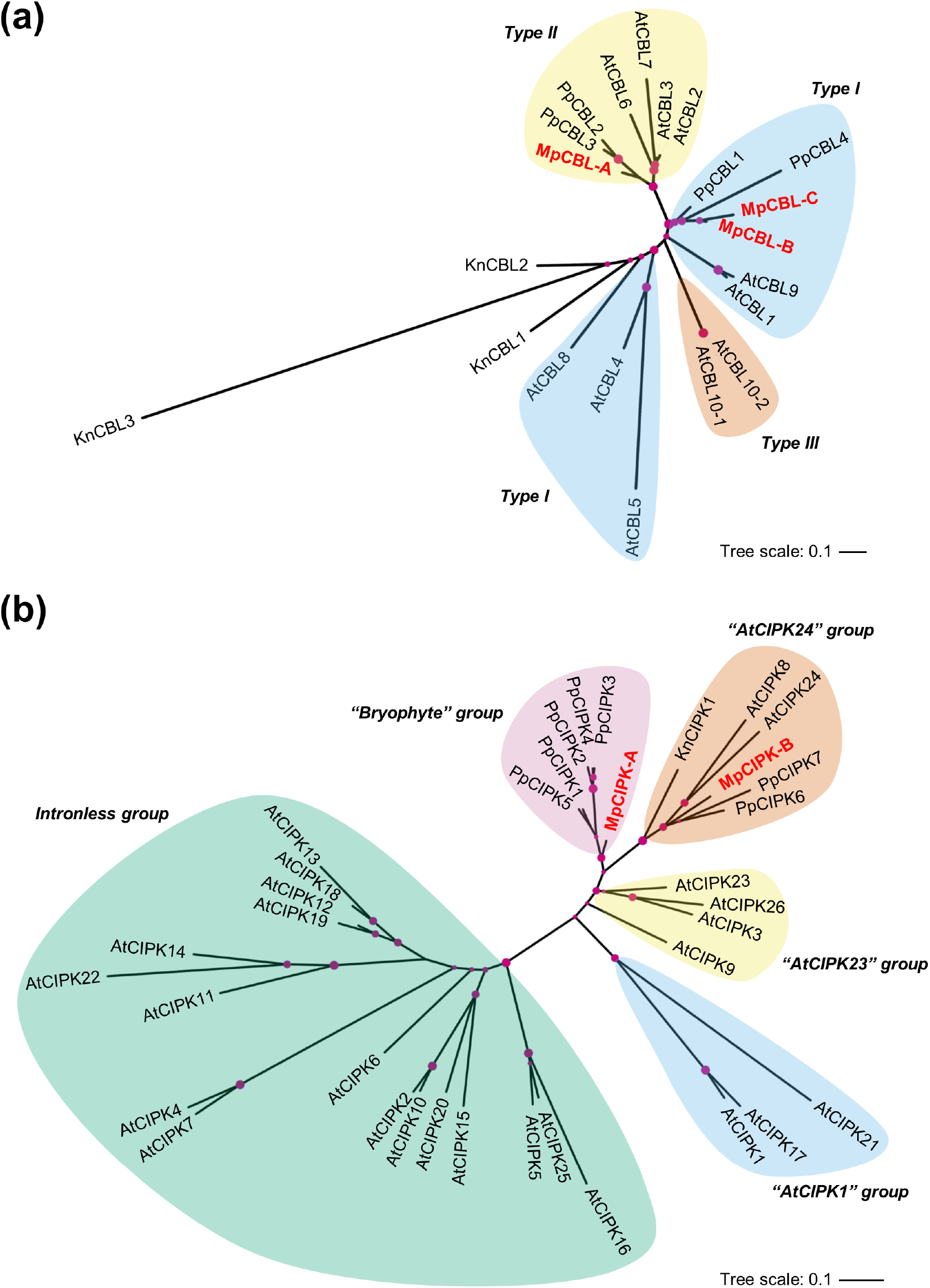
Marchantia polymorpha has three CBLs and two CIPKs. Unrooted phylogenetic trees show sequence relatedness of CBL (a) and CIPK (b) proteins from Arabidopsis thaliana (At), Physcomitrium patens (Pp), Klebsormidium nitens (Kn) and Marchantia polymorpha (Mp; red text). Shading denotes CBL types and CIPK groups according to classifications in Edel & Kudla (2015). Pink circles represent 80-100% bootstrapping support for each branch by size.

Two MpCIPKs were identified in version 5.0 of the *M. polymorpha* genome (Fig. 1b). Two clades of CIPKs are known and defined based on intron number, with the intronless clade seemingly arising in angiosperms (Zhu *et al*., 2016). Both MpCIPKs fall into the intron-rich clade of CIPKs, as would therefore be expected, and both MpCIPKs contain the NAF domain that is both necessary and sufficient for CBL-CIPK interactions (Albrecht *et al*., 2001). Seven other potential MpCIPKs were identified but excluded based on the lack of a detectable NAF domain. MpCIPK-A falls into the “bryophyte” group with PpCIPK1/2/3/4/5, which seems to represent a group that diverged in bryophytes and has no representatives from *A. thaliana* or *Klebsormidium nitens* (Fig. 1b). MpCIPK-B is similar to PpCIPK7 from *P. patens* and falls into the “AtCIPK24” group, which harbours AtCIPK8 and AtCIPK24 (SOS2). *K. nitens* has a single CIPK that falls into the “AtCIPK24” group which may imply that this is the earliest clade of CIPKs present in land plants (Kleist *et al*., 2014). Both AtCIPK8/24 and PpCIPK1 operate in salt sensitivity pathways (Quan *et al*., 2007; Yin *et al*., 2020; Xiao *et al*., 2021), implying that salt stress signalling might be the original function of CIPKs in the “AtCIPK24” group and in fact the main function of the CIPKs in *M. polymorpha*.

### Different *Marchantia polymorpha* accessions show similar salt stress tolerance responses

To establish whether common salt tolerance mechanisms exist in *M. polymorpha*, we compared the salt stress responses of different plant accessions. Two *M. polymorpha* accessions from different regions have been widely used in research to date: Tak-1/2 from Takaragaike Park in Kyoto (Japan) and Cam-1/2 from Cambridge (United Kingdom). Utilising common nomenclature, male plants are labelled 1 (e.g. Tak-1) and female plants are labelled 2 (e.g. Tak-2). Male and female plants of both accessions were grown in the presence of different concentrations of salt for one week, after which growth was assessed and total fresh weight was measured. All plants survived treatments of 150 mM NaCl for one week and decreased growth was observed in response to increasing salt concentration (Fig. 2a, b). Treatments with 100 and 150 mM NaCl caused significant decreases in fresh weight relative to control plants, but treatments with 50 mM NaCl did not cause a significant decrease (Fig. 2b). Since statistical differences in fresh weight were observed between thallus tissue of the same size from Tak-1 and Cam-1 plants, normalised fresh weight was used to compare between accessions (Fig. 2b). These comparisons reveal that Tak-1, Tak-2, Cam-1 and Cam-2 plants all demonstrate similar salt tolerance responses from 0-150 mM salt, with no significant differences observed between male and female plants or between the Takaragaike and Cambridge accessions (Fig. 2b).

**Figure 2:**
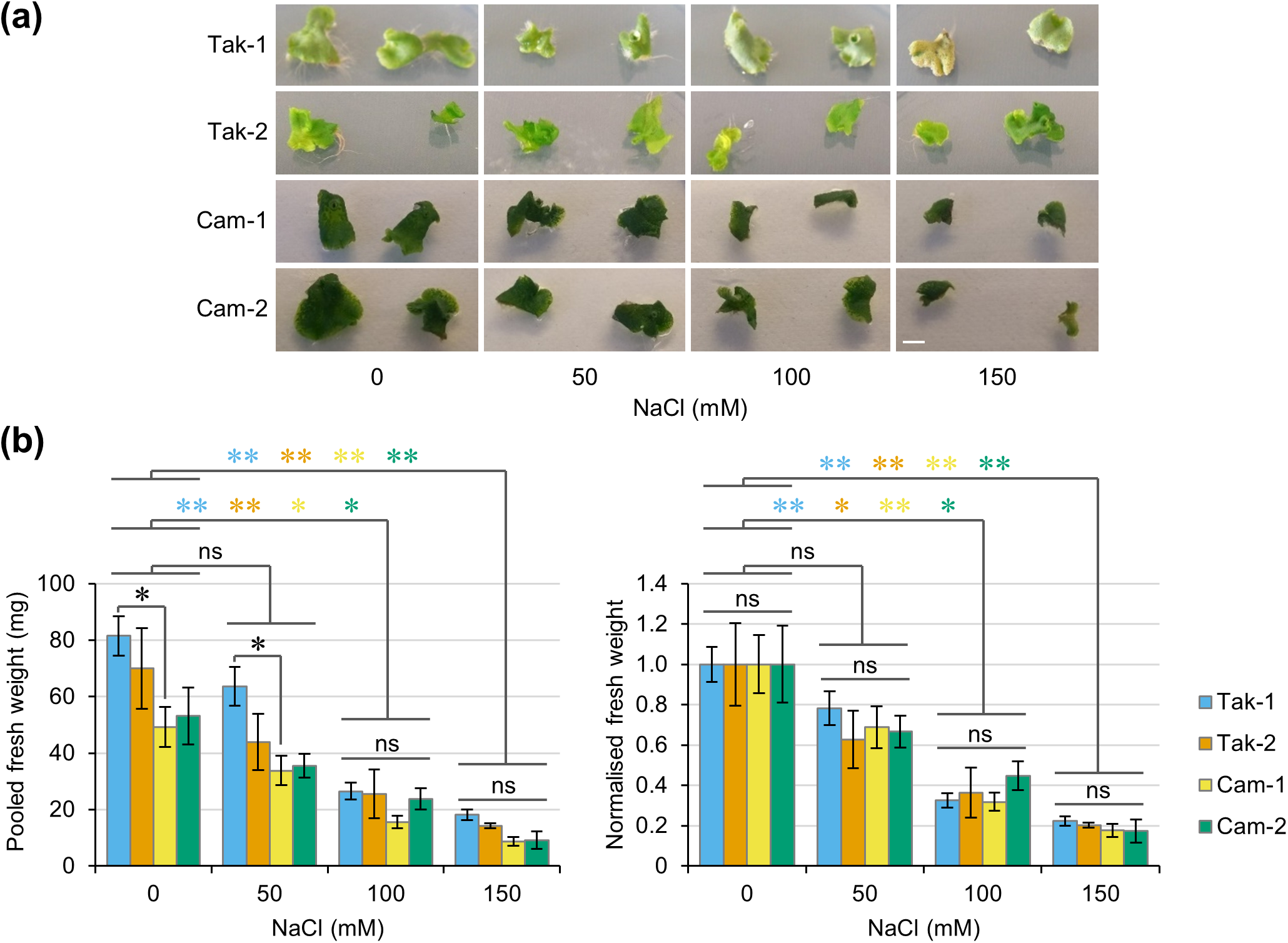
Different accessions of Marchantia polymorpha show no difference in salt stress tolerance. (a) Representative images of thallus tissue from Tak-1, Tak-2, Cam-1 and Cam-2 plants grown in the presence of the indicated concentrations of NaCl for 7 days. (b) Graphs show pooled fresh weight and normalised fresh weight (normalised to samples of the same genotype grown with 0 mM NaCl) of 5 pieces of thallus tissue from (a). Data represent mean ± standard error from at least four independent biological replicates. Significance in an ANOVA with post-hoc Tukey test is indicated at p<0.05 (one star) or p<0.01 (two stars). Black stars denote significant difference between accessions grown under the same condition, while coloured stars denote significance between treatments for the indicated accessions. Scale bar = 5 mm; ns = not significant.

### *CIPK-B* expression is regulated by salt stress

To determine if any of the *M. polymorpha* CIPKs and CBLs function in a salt sensitivity pathway, expression of the *MpCBLs* and *MpCIPKs* was investigated in plants grown for one week in the presence of 0-150 mM NaCl. In *A. thaliana*, the SOS pathway is upregulated in the first 24 hours of salt stress but then downregulated after day six (Ji *et al*., 2013; Rolly *et al*., 2020). We therefore expected early salt-responsive *CBLs* and *CIPKs* in *M. polymorpha* to also show downregulated expression after a salt stress treatment of one week. Cam-2 (Fig. 3) and Tak-1 (Fig. S1) plants showed strong upregulation of the abiotic stress marker gene *LEA-like4* upon salt treatments and both accessions had salt-responsive *MpCIPKs* and *MpCBLs*. In Cam-2, both *MpCIPK-A/B* were downregulated in response to salt treatments from 100-150 mM NaCl (Fig. 3a). Similar downregulation of *MpCIPK-A/B* was also observed in Tak-1 at 50 mM NaCl (Fig. S1). Interestingly, in both accessions, downregulation of *CIPK-B* was observed in a greater number of salt stress treatments than *CIPK-A*. Therefore, *MpCIPK-A/B* likely maintain functions in either salt- or drought-responsive pathways, with *CIPK-B* possibly more important in these processes. *MpCBL-C* was downregulated in response to all salt stress treatments in Cam-2 (Fig. 3a), and *MpCBL-C* was also downregulated in Tak-1 under 50 and 100 mM NaCl treatments (Fig. S1). *MpCBL-A/B* showed some downregulation in response to salt stress, but this was more modest than *MpCBL-C* downregulation and somewhat variable between accessions. The consistent and strong downregulation of *MpCBL-C* expression in both accessions implies that *MpCBL-C* may function in salt- or drought-responsive tolerance in *M. polymorpha* more widely. In contrast, changes in *MpCBL-A/B* expression were not consistent between the two accessions and therefore *MpCBL-A/B* may have subtly different functions or regulation between the accessions.

**Figure 3:**
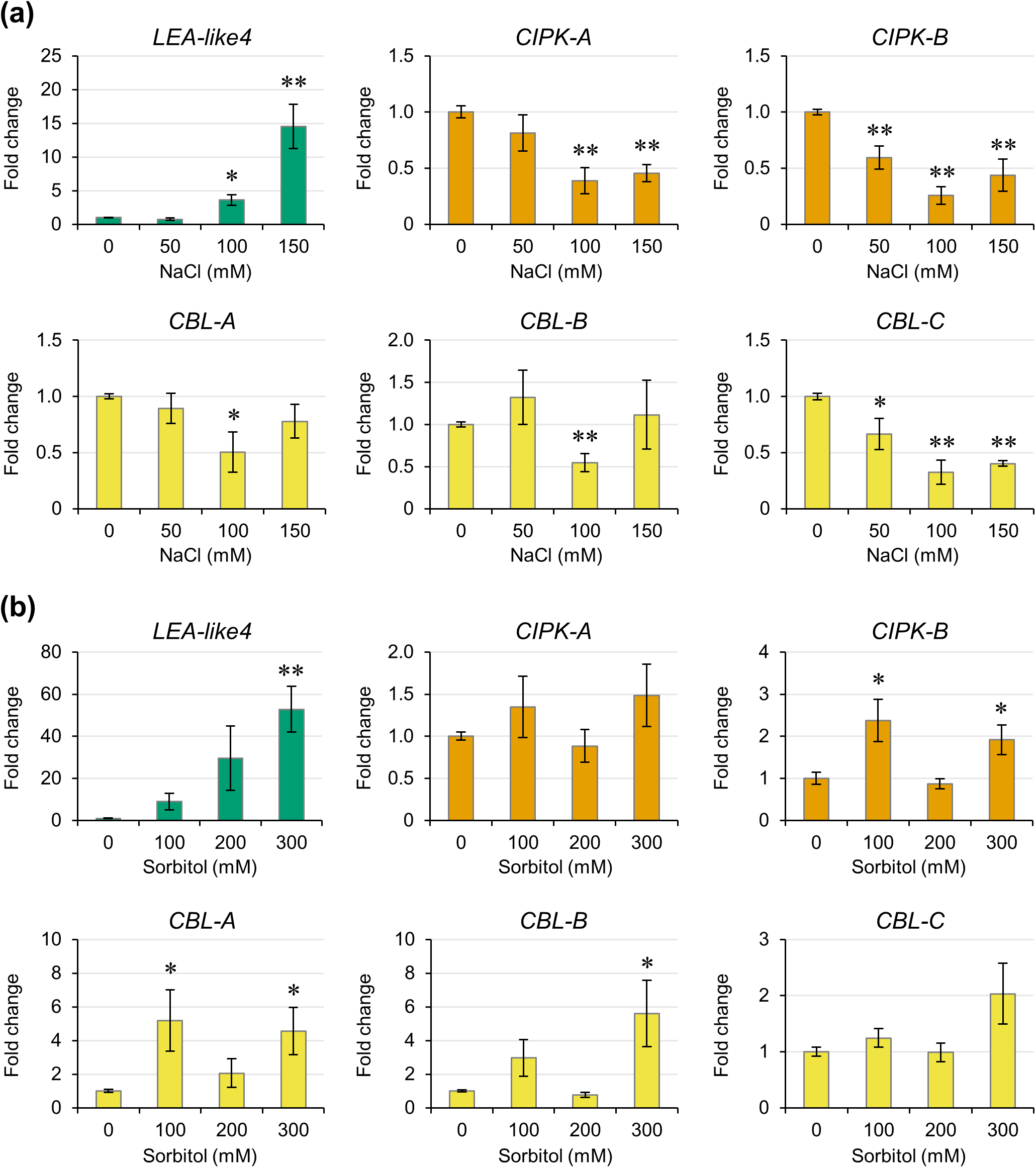
Expression of CIPK-B is downregulated by salt stress but upregulated by osmotic stress. Expression changes of the indicated genes were measured by RT-qPCR in thallus tissue of Cam-2 plants grown in the presence of the indicated concentrations of NaCl (a) or sorbitol (b) for 7 days. Significance in a pairwise two-tailed t-test relative to 0 mM treatment is indicated at p<0.05 (one star) or p<0.01 (two stars). Data represent mean ± standard error from three independent biological replicates.

As *MpCBL-C* and *MpCIPK-B* show transcriptional changes in two accessions grown under salt stress, it is most likely that these are the salt-responsive CBL and CIPK in *M. polymorpha*. However, these transcriptional responses could also be caused by the drought component of salt stress. To determine if *CBL-C* and *CIPK-B* therefore play roles in osmotic stress, we grew Cam-2 plants for one week in the presence of isoosmolar concentrations of the osmoticum sorbitol and assessed gene expression via RT-qPCR. The abiotic stress marker gene *LEA-like4* was again strongly induced by growth on treatment plates (Fig. 3b). *MpCIPK-A* showed no significant changes in gene expression, but *MpCIPK-B* showed significant upregulation (Fig. 3b). Of the CBLs, *MpCBL-A/B* both showed significant upregulation, whilst *MpCBL-C* expression was unchanged with sorbitol (Fig. 3b). This suggests *MpCBL-A/B* may play roles in osmotic stress, while the likely function of *MpCBL-C* in salt stress may be due to ionic rather than osmotic stress. Overall, this data shows that the transcriptional responses of *CIPK-B* and *CBL-C* are different upon salt stress and osmotic stress, and that the downregulation of these genes during salt stress is specific to ionic stress.

### CBLs and CIPKs from *Marchantia polymorpha* do not form specific protein-protein interactions

Protein-protein interactions between different CBLs and CIPKs were assessed, including AtSOS3 (AtCBL4) and AtSOS2 (AtCIPK24), to determine which MpCBLs and MpCIPKs can interact and whether any of the MpCBLs and MpCIPKs may constitute a SOS pathway in *M. polymorpha*. Yeast two-hybrid assays revealed that MpCIPK-A could interact with all MpCBLs and AtCBL4, while MpCIPK-B interacted with CBL-A and CBL-B but not CBL-C or AtCBL4 (Fig. 4a; Fig. S2). AtCIPK24 (SOS2) specifically interacted with CBL-B and CBL-C. Removal of the NAF domain abolished all CBL-CIPK interactions (Fig. 4a), confirming that *M. polymorpha* CBL-CIPK interactions occur in an equivalent manner to *A. thaliana*, where the NAF domain is necessary and sufficient for CBL-CIPK interactions (Albrecht *et al*., 2001). Based on these protein-protein interactions, we would expect that CBL-C or CBL-B could be AtCBL4 homologues. Moreover, MpCIPK-A could be a potential SOS2 homologue as it can interact with AtCBL4. However, since MpCIPK-A interacts with all MpCBLs and AtCBL4, it may be that MpCIPK-A simply forms non-specific interactions in the yeast system. MpCBL-B also interacts with both MpCIPKs and AtCIPK24, and may therefore also be non-specific for the interactions it forms. The similar protein-protein interactions observed for MpCBL-C and AtCBL4, alongside the *MpCBL-C* expression data, support the hypothesis that MpCBL-C may be a salt sensitive AtSOS3 homologue. However, it is notable that CIPK-B and CBL-C do not interact in our yeast two-hybrid assays. This appears to contradict our hypothesis that CIPK-B and CBL-C function in salt stress signalling, so we sought to validate all protein-protein interactions assessed by yeast two-hybrid using additional *in planta* assays.

**Figure 4:**
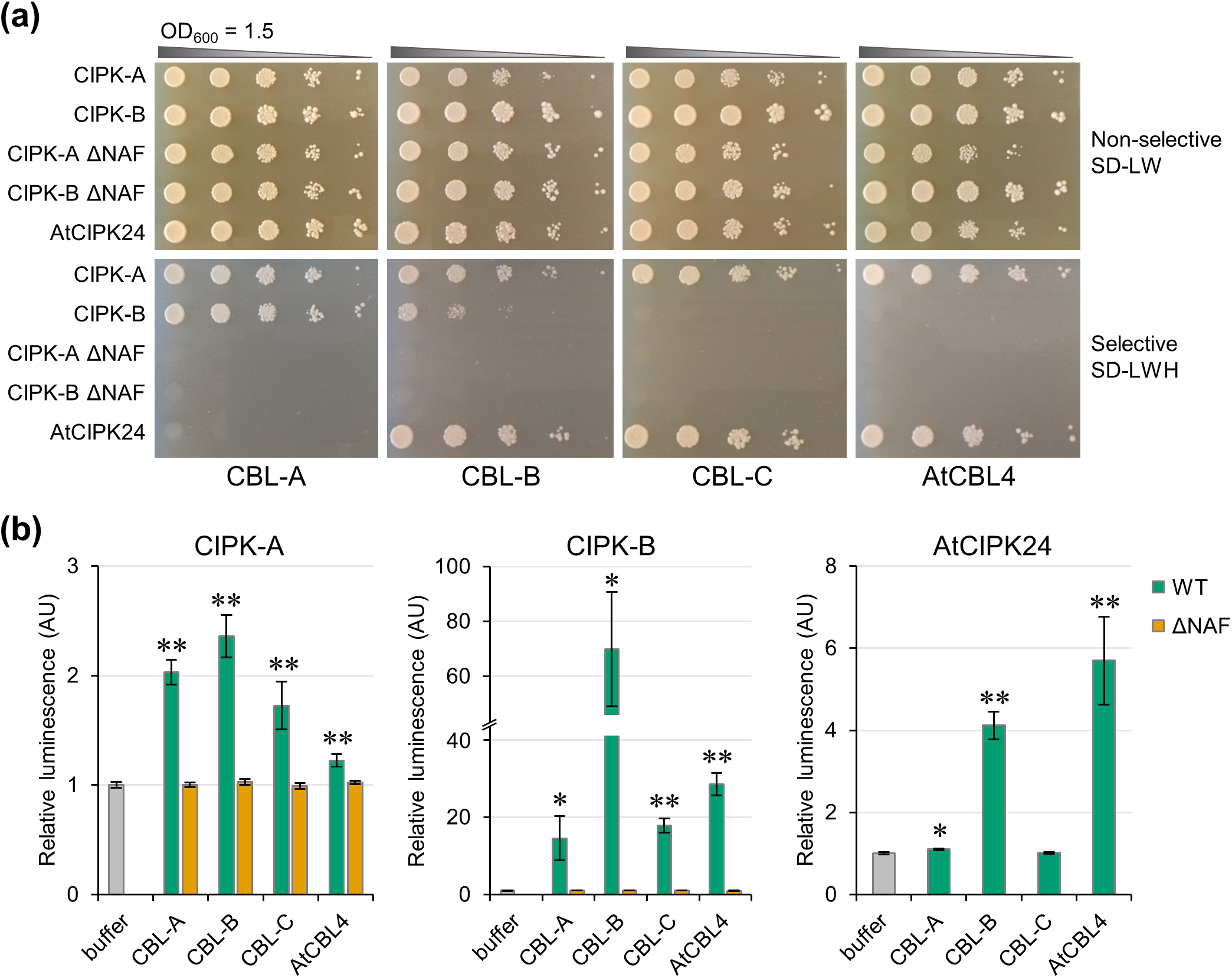
CIPK-B forms protein-protein interactions with all *M. polymorpha* CBLs and Arabidopsis thaliana CBL4. (a) Representative images of yeast two-hybrid assay results, showing serial dilution of yeast cells grown on non-selective (SD-Leu-Trp; SD-LW) or selective media (SD-Leu-Trp-His; SD-LWH) as indicated. All interactions were tested on at least three independent occasions, with identical results obtained for each replicate. (b) Split luciferase assays were performed in N. benthamiana to test CBL-CIPK interactions as indicated. Significance in a pairwise two-tailed t-test relative to buffer control is indicated at p<0.05 (one star) or p<0.01 (two stars). Data represent mean ± standard error from three independent biological replicates.

To confirm the results from the yeast two-hybrid system, a split luciferase assay was performed to determine the interactions formed between MpCBLs and MpCIPKs in *N. benthamiana*. Importantly, all interactions between MpCBLs and MpCIPKs were detectable in this *N. benthamiana* system (Fig. 4b), and all interactions were dependent on the presence of the NAF domain. AtCIPK24 (SOS2) formed interactions with MpCBL-*A* and MpCBL-B, but not MpCBL-C (Fig. 4b). Bimolecular fluorescence complementation assays were also performed in *N. benthamiana* leaves, confirming interactions between all MpCBLs and MpCIPKs (Fig. S3). Since the results of the two *in planta* assays support each other and identified interactions that could not be detected in the yeast two-hybrid assays, we conclude that MpCBLs do not form specific protein-protein interactions with MpCIPKs. Therefore all CBL-CIPK interactions are likely possible in *M. polymorpha*.

### CIPK-B is involved in tolerance to salt stress but not osmotic stress

Two independent knockout lines of *CIPK-B* from *M. polymorpha* were generated using the CRISPR/Cas9 system (Fig. 5a). Both *cipk-b* mutant lines were smaller than wildtype (Cam-2) plants (Fig. 5b), suggesting a possible role for *CIPK-B* in plant growth and development. To test whether *cipk-b* mutants were impaired in responses to salt stress, equal-sized pieces of thallus tissue were excised from mature plants, grown in the presence of different concentrations of NaCl for one week, and the fresh weight was measured. To account for any differences in the starting weight between wildtype and *cipk-b* mutant plants, the fresh weight for each line was also normalised to growth on control plates (0 mM NaCl). Increased salt in the growth media resulted in decreased growth of wildtype plants and both *cipk-b* mutants (Fig. 5c, d). However, both *cipk-b* mutant lines demonstrated reduced salt tolerance, showing decreased growth and biomass under mild salt stress (50 mM NaCl) compared with wildtype plants (Fig. 5c, d). Both *cipk-b* mutants also showed increased chlorosis and death from as low as 50 mM NaCl compared with wildtype plants, which showed tolerance up to 150 mM NaCl (Fig. 5c). Chlorophyll content was lower in both *cipk-b* mutant lines compared with wildtype plants under all salt stress conditions (Fig. 5e).

**Figure 5:**
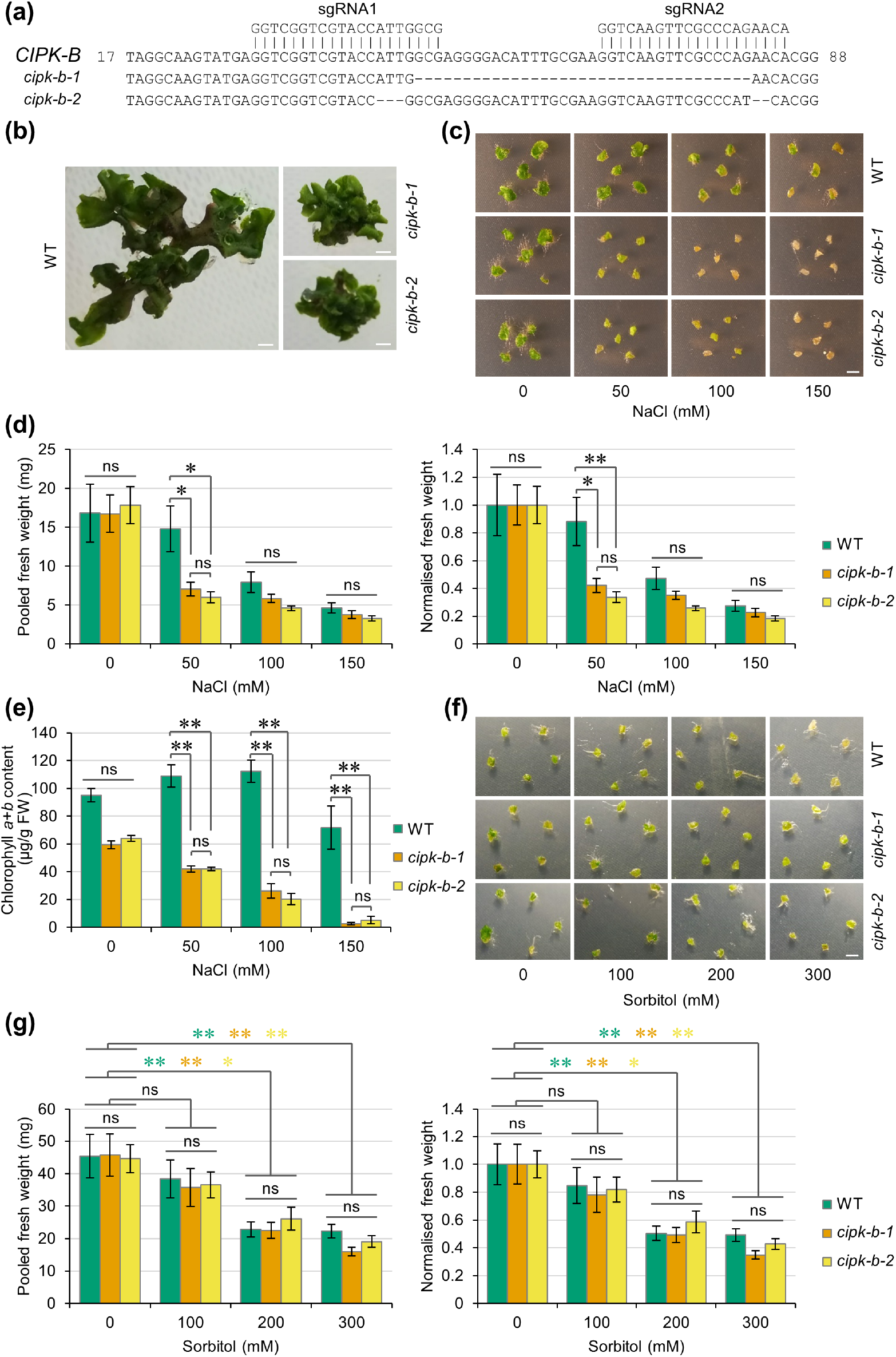
*cipk-b* mutants are impaired for tolerance to salt stress but not osmotic stress. (a) Two 20-nucleotide sgRNAs (indicated) were designed to target exon 1 of CIPK-B to generate two independent cipk-b mutant lines via CRISPR/ Cas9 gene editing in the Cam-2 background. The cipk-b-1 mutant contains a 35 bp deletion and the cipk-b-2 mutant contains two small deletions and a single nucleotide substitution (as indicated), both leading to missense mutations and introduction of a premature stop codon (resulting in predicted proteins containing 29 and 39 amino acids in the cipk-b-1 and cipk-b-2 mutant lines, respectively). (b) Representative images of size and morphological differences between WT (Cam-2) and both cipk-b mutant lines grown from gemmae for six weeks in the absence of salt stress. (c) Representative images of thallus tissue from WT (Cam-2) and both cipk-b mutant lines grown in the presence of the indicated concentrations of NaCl for 7 days. (d) Graphs show pooled fresh weight and normalised fresh weight (normalised to samples of the same genotype grown with 0 mM NaCl) of 5 pieces of thallus tissue from (c). Data represent mean ± standard error from at least five independent biological replicates. Significance in a Dunn’s test is indicated at p<0.05 (one star) or p<0.01 (two stars). (e) Graph shows chlorophyll a+b content in WT (Cam-2) and both cipk-b mutant lines in material from (c). Data represent mean ± standard error from three independent biological replicates. Significance in a Dunn’s test is indicated at p<0.05 (one star) or p<0.01 (two stars). (f) Representative images of thallus tissue from WT (Cam-2) and both cipk-b mutant lines grown in the presence of the indicated concentrations of sorbitol for 7 days, which are isoosmolar to the NaCl concentrations used in (c). (g) Graphs show pooled fresh weight and normalised fresh weight (normalised to samples of the same genotype grown with 0 mM sorbitol) of 5 pieces of thallus tissue from (f). Data represent mean ± standard error from at least five independent biological replicates. Significance in a Dunn’s test is indicated at p<0.05 (one star) or p<0.01 (two stars). Coloured stars denote significance between treatments for the indicated plant lines. Scale bar in all panels = 5 mm; ns = not significant.

To test if CIPK-B is involved in the ionic or osmotic component of salt stress, treatments were also performed using isoosmolar concentrations of sorbitol. Wildtype and both *cipk-b* mutant lines showed decreased growth in response to increasing concentrations of sorbitol (Fig. 5f, g), but no significant difference was observed in the pooled or normalised fresh weight between wildtype plants and either *cipk-b* mutant. We therefore conclude that CIPK-B does not have a role in osmotic stress and that the salt stress phenotype observed for *cipk-b* mutant plants is rather due to ionic responses associated with salt stress.

## DISCUSSION

A number of distinct signalling pathways have been characterised in *M. polymorpha*, including those using auxin, cytokinin, jasmonate, and ethylene. Signalling in these pathways is similar to that in angiosperms, but with fewer pathway components or instead utilising precursors of the hormone. Many of these reduced signalling pathways may represent signalling present in the last common ancestor between bryophytes and other land plants. However, there is some evidence of reductive evolution in *M. polymorpha*, including stomata loss and subsequent development of air pores (Harris *et al*., 2020). Therefore, to have confidence in inferring the state of the last universal common ancestor of land plants, multiple bryophytes need to be interrogated. This is also true for understanding calcium signalling in bryophytes. To date, two studies have investigated *P. patens* (Kleist *et al*., 2014; Xiao *et al*., 2021), but, as far as we are aware, this is the first study of CBL-CIPK calcium decoders in *M. polymorpha*. Our work demonstrates a clear role for MpCIPK-B in salt stress signalling. MpCIPK-B falls into the same clade as AtCIPK24 (SOS2; Fig. 1b) and *cipk-b* knockout mutants showed increased salt sensitivity (Fig. 5). Since *cipk-b* mutants did not respond to sorbitol of isoosmolar concentrations, we conclude that MpCIPK-B is ionic-specific and not drought-responsive more generally.

Previously, the *P. patens* genome was interrogated and six PpCBLs and eight PpCIPKs were identified; however, two of these CBLs and one CIPK showed no evidence of being expressed or were pseudogenes (Kleist *et al*., 2014). Utilising version 5.0 of the *M. polymorpha* genome, we identified three CBLs and two CIPKs. MpCBL-B/C were in the Type I CBL group with identifiable MGCxxS/T palmitoylation and myristoylation motifs for plasma membrane targeting, with close grouping to PpCBL4 in a branch containing AtCBL1/9 and PpCBL1. MpCBL-A fell into the Type II CBL group with no detectable motif for tonoplast targeting, defined as multiple cysteine residues in the first 20 amino acids for S-acylation (Batistič *et al*., 2012; Tang *et al*., 2012). The closest related CBLs to MpCBL-A include PpCBL2/3 and AtCBL2/3/6/7, which are known to localise to the tonoplast. Whether MpCBL-A localises to the tonoplast remains to be determined, but this may be interesting to explore further given that MpCBL-A lacks a canonical tonoplast targeting motif and subcellular targeting motifs in *M. polymorpha* may be different from angiosperms. No Type III CBLs were identified in this study or previously for PpCBLs or MpCBLs (Kleist *et al*., 2014; Edel & Kudla, 2015) indicating that this group of CBLs may not be present in bryophytes, or was subsequently lost by both *P. patens* and *M. polymorpha*.

CIPKs in the last universal common ancestor of land plants were likely to be intron-rich and subsequent evolution has led to much wider diversification. Two MpCIPKs were identified here and in a previous study (Edel & Kudla, 2015). Of these, MpCIPK-A falls into a clade of bryophyte CIPKs alongside PpCIPK1/2/3/4/5 and likely represents early signalling functions of CIPKs (Kleist *et al*., 2014). MpCIPK-B falls into a group including PpCIPK6/7 and AtCIPK8/24, which likely represents an early group that arose with progression onto land. AtCIPK24 (SOS2) is known to confer salt tolerance through phosphorylation-based activation of SOS1, a Na^+^/H^+^ antiporter downstream of calcium decoding by SOS3 (AtCBL4) or ScaBP8 (AtCBL10). AtCIPK8 has also been characterised in salt tolerance through the same pathway through interaction with ScaBP8 (AtCBL10). In addition to MpCIPK-B, the only other characterised bryophyte CBL or CIPK involved in salt tolerance of which we are aware is PpCIPK1 (Xiao *et al*., 2021). Interestingly, PpCIPK1 falls into the bryophyte CIPK clade (with MpCIPK-A) and not the AtCIPK24 clade (with MpCIPK-B; Fig. 1b), suggesting that the general function of several of the early land plant CIPKs may also be in response to salt tolerance.

Previous studies in *A. thaliana* have investigated *CBL* and *CIPK* expression after 24 hours of salt treatment and demonstrated that components of the SOS pathway are upregulated within 24 hours but then downregulated at later time points (Ji *et al*., 2013; Rolly *et al*., 2020). A number of CBLs and CIPKs are involved in ionic balance in angiosperms (such as AtCBL1/9 and AtCIPK23; Li *et al*., 2006; Cheong *et al*., 2007; Ragel *et al*., 2015) and these too are upregulated within 24 hours, despite not being directly involved in Na^+^ tolerance but rather in Na^+^/K^+^ equipoise. We have analysed a previously published RNA sequencing dataset and confirmed that all *MpCBLs* are upregulated within 24 hours of salt stress and that neither *MpCIPK* shows changes in expression within this time frame (Wu *et al*., 2021). As our RT-qPCR confirms downregulation of *CBL-C, CIPK-A* and *CIPK-B* in both the Cam-2 and Tak-1 accessions after one week of salt stress (Fig. 3; Fig. S1), we are confident that either or both MpCIPKs have some involvement in drought and salt tolerance responses alongside MpCBL-C.

The finding that all MpCBLs and MpCIPKs interact *in planta* is important. Our *M. polymorpha* finding is different from studies on PpCBLs and PpCIPKs, where specific protein-protein interactions have been identified (Xiao *et al*., 2021), and suggests that either specificity was lost or greater versatility of the CBL-CIPK network arose following whole genome duplication. Many previous studies have utilised yeast two-hybrid to test CBL-CIPK interactions and then confirmed positive protein-protein interactions *in planta*. From our data, it is evident that such an approach may fail to detect important CBL-CIPK interactions due to the prominence of false negative results with the yeast two-hybrid method. For instance, yeast two-hybrid assays revealed that MpCIPK-B could not interact with AtCBL4 and MpCBL-C (Fig. 4a), which we believe to be the main salt signalling *M. polymorpha* CBL based on phylogeny (Fig. 1a) and gene expression (Fig. 3; Fig. S1), but positive interactions were detected between these proteins when tested *in planta* (Fig. 4b; Fig. S3). The comparatively small CBL-CIPK network of *M. polymorpha* means that we have been able to analyse carefully *in planta* the protein-protein interactions formed by an entire CBL-CIPK network. This important analysis reveals a surprising lack of specificity for CBL-CIPK interactions in *M. polymorpha*. This lack of specificity may not necessarily mean that all CBL-CIPK combinations are capable of forming in a single cell at the same time, to transduce different calcium signals into kinase activity and substrate phosphorylation. For example, some CBLs or CIPKs may show different spatial or temporal expression profiles, but more work is needed in the future to explore this further. It would also be interesting to investigate *P. patens* and *M. polymorpha* CBL-CIPK interactions further to determine how the specificity of these interactions arose, and if *M. polymorpha* CBLs can interact with *P. patens* CIPKs (or vice versa). Such work may also determine whether evolution of specificity of the protein-protein interaction was primarily driven by the CBL or the CIPK. Previous structural work implies that specificity of CBL-CIPK interactions comes from the tertiary structure of the CBL, either allowing or disallowing interactions (Sánchez-Barrena *et al*., 2005); however, it is still unknown how phosphorylation causes structural changes that can facilitate interactions, for example as in AtCBL10 (Xie *et al*., 2009).

Together, our work suggests that the general function of several of the early land plant CIPKs is in salt tolerance responses. We suggest that additional functions to regulate other ionic stresses may then have evolved later and possibly as a result of genome duplication that gave rise to subsequent expansion and increased versatility of the CBL-CIPK network. Additional tests are required to interrogate these early groups of CIPKs and CBLs to understand fully the original functions of calcium signalling and decoder proteins in the evolution of land plants. Further investigation of other calcium decoder families will give understanding of the evolution of unique calcium decoder function, and how neofunctionalization occurs in a complex system of encoding, decoding and transducing calcium signals to give rise to tolerance responses. Modulating these signalling processes in plants will be key to future solutions to engineer tolerance responses in crops in the face of global problems of salinization in agriculture.

## ACKNOWLEDGEMENTS

We thank Jim Haseloff for providing wildtype plant material, and Susana Sauret-Gueto and Linda Silvestri for transformation tips for *M. polymorpha*. We also thank Nico Pelaez Llaneza for assistance with fluorescence microscopy. CT was funded by a PhD studentship from the Biotechnology and Biological Sciences Research Council (BBSRC) via the Norwich Research Park Doctoral Training Partnership (NRPDTP). BW was funded by a BBSRC Research Experience Placement (REP) grant. JBM was supported by an Eastern ARC fellowship. RDP and JBM were funded by the Natural Environment Research Council (NE/V000756/1) and BBSRC (BB/X005968/1).

## AUTHOR CONTRIBUTIONS

CT, TW and JBM conceptualised the study. CT and TW carried out the phylogenetic analyses; CT performed phenotyping of wildtype plants (Takaragaike and Cambridge accessions) grown under salt stress; CT and AMER carried out gene expression analyses by RT-qPCR during salt and osmotic stress, respectively; CT performed yeast two-hybrid analysis, western blotting and BiFC assays; JH carried out split luciferase assays; CT performed CRISPR/Cas9 knockout mutant generation; CT and JH performed phenotyping of wildtype and *cipk-b* mutant lines grown under salt stress; JH and RDP carried out the chlorophyll measurements; AMER and BW performed phenotyping of wildtype and *cipk-b* mutant lines grown under osmotic stress. CT and JBM wrote the manuscript, with input and critical feedback from all co-authors.

## DATA AVAILABILITY

The data that support the findings of this study are available from the corresponding author upon reasonable request.

## Notes

### Competing Interest Statement

The authors have declared no competing interest.

