## Supplemental Tables for "CIPK-B is essential for salt stress signalling in *Marchantia polymorpha*"

### SUPPORTING INFORMATION

**Table S1: Gene ID numbers for sequences used in bioinformatics in this study**

| Gene name | Species | Gene ID |
|---|---|---|
| *MpCBL-A* | *M. polymorpha* | Mp2g07750 (Mapoly0015s0061) |
| *MpCBL-B* | *M. polymorpha* | Mp4g00900 (Mapoly0066s0053) |
| *MpCBL-C* | *M. polymorpha* | Mp5g19810 (Mapoly0134s0040) |
| *MpCIPK-A* | *M. polymorpha* | Mp1g05680 (Mapoly0005s0039) |
| *MpCIPK-B* | *M. polymorpha* | Mp2g26670 (Mapoly0025s0017) |
| *KnCBL1* | *K. nitens* | GAQ81593.1 |
| *KnCBL2* | *K. nitens* | GAQ84394.1 |
| *KnCBL3* | *K. nitens* | GAQ80563.1 |
| *KnCIPK1* | *K. nitens* | GAQ84395.1 |
| *PpCBL1* | *P. patens* | Pp3c1_36780V3.1.p |
| *PpCBL2* | *P. patens* | Pp3c16_24350V3.1.p |
| *PpCBL3* | *P. patens* | Pp3c13_3530V3.1.p |
| *PpCBL4* | *P. patens* | Pp3c5_9970V3.1.p |
| *PpCIPK1* | *P. patens* | Pp3c16_15230V3.1.p |
| *PpCIPK2* | *P. patens* | Pp3c2_13790V3.1.p |
| *PpCIPK3* | *P. patens* | Pp3c10_7160V3.1.p |
| *PpCIPK4* | *P. patens* | Pp3c14_7960V3.1.p |
| *PpCIPK5* | *P. patens* | Pp3c5_7750V3.1.p |
| *PpCIPK6* | *P. patens* | Pp3c12_210V3.1.p |
| *PpCIPK7* | *P. patens* | Pp3c15_1210V3.1.p |

**Table S2: Primers used for RT-qPCR in this study**

| Gene name | Gene ID | Forward primer | Reverse primer |
|---|---|---|---|
| *CBL-A* | Mp2g07750 | AGCGGAAAGAGGTGAAACGG | GAGAGGGATGCTGCTGAACC |
| *CBL-B* | Mp4g00900 | GGGCTGCTTCAGCTCAAAAC | CGCAAGCTGGAACTCTTCCT |
| *CBL-C* | Mp5g19810 | CAAGTGCTCCACCAGAGGAC | GCCTCCGCAAATGTCTTGTC |
| *CIPK-A* | Mp1g05680 | AAACACCCTGCGAACGAGAT | ACCTCAAACACCTCTGTGGC |
| *CIPK-B* | Mp2g26670 | CCTGTACGGATGCACGATGA | AGAACGGAAAGGTTGAGCCC |
| *LEA-like4* | Mp1g23200 | GCTAACAGACCCAGGTGAC | TGTTTCCAACGGCAGAGTG |
| *ACT1* | Mp6g10990 | GAGCGCGGTTACTCTTTCAC | GACCGTCAGGAAGCTCGTAG |
| *APT* | Mp3g35140 | CGAAAGCCCAAGAAGCTACC | GTACCCCCGGTTGCAATAAG |

**Table S3: Final constructs created and used in this study**

| ID | Construct details | Experiment |
|---|---|---|
| 01044 | pTRP1-LEU2-tADH1 pADH1-GAL4-AD-3xHA-MpCIPK-A-tADH1 | Yeast two-hybrid GAL4-AD-MpCIPK-A |
| 01045 | pTRP1-LEU2-tADH1 pADH1-GAL4-AD-3xHA-MpCIPK-B-tADH1 | Yeast two-hybrid GAL4-AD-MpCIPK-B |
| 01046 | pTRP1-TRP1-tADH1 pADH1-GAL4-BD-3xc-myc-MpCBL-A-tADH1 | Yeast two-hybrid GAL4-BD-MpCBL-A |
| 01147 | pTRP1-TRP1-tADH1 pADH1-GAL4-BD-3xc-myc-MpCBL-B-tADH1 | Yeast two-hybrid GAL4-BD-MpCBL-B |

| | | |
|---|---|---|
| 01048 | pTRP1-TRP1-tADH1<br>pADH1-GAL4-BD-3xc-myc-MpCBL-C-tADH1 | Yeast two-hybrid<br>GAL4-BD-MpCBL-C |
| 01133 | pTRP1-LEU2-tADH1<br>pADH1-GAL4-AD-3xHA-MpCIPK-A-ΔNAF-tADH1 | Yeast two-hybrid<br>GAL4-AD-MpCIPK-A<br>ΔNAF |
| 01134 | pTRP1-LEU2-tADH1<br>pADH1-GAL4-AD-3xHA-MpCIPK-B-ΔNAF-tADH1 | Yeast two-hybrid<br>GAL4-AD-MpCIPK-B<br>ΔNAF |
| 01145 | pTRP1-TRP1-tADH1<br>pADH1-GAL4-BD-3xc-myc-AtCBL4-tADH1 | Yeast two-hybrid<br>GAL4-BD-AtCBL4 |
| 01146 | pTRP1-LEU2-tADH1<br>pADH1-GAL4-AD-3xHA-AtCIPK24-tADH1 | Yeast two-hybrid<br>GAL4-AD-AtCIPK24 |
| 01268 | p35S-MpCBL-A-LgBiT-t35S<br>pNOS-SmBiT-MpCIPK-A-tNOS<br>pAtUBI10-GUS-tNOS | Split luciferase<br>MpCBL-A-LgBiT<br>SmBiT-MpCIPK-A |
| 01269 | p35S-MpCBL-B-LgBiT-t35S<br>pNOS-SmBiT-MpCIPK-A-tNOS<br>pAtUBI10-GUS-tNOS | Split luciferase<br>MpCBL-B-LgBiT<br>SmBiT-MpCIPK-A |
| 01270 | p35S-MpCBL-C-LgBiT-t35S<br>pNOS-SmBiT-MpCIPK-A-tNOS<br>pAtUBI10-GUS-tNOS | Split luciferase<br>MpCBL-C-LgBiT<br>SmBiT-MpCIPK-A |
| 01271 | p35S-AtCBL4-LgBiT-t35S<br>pNOS-SmBiT-MpCIPK-A-tNOS<br>pAtUBI10-GUS-tNOS | Split luciferase<br>AtCBL4-LgBiT<br>SmBiT-MpCIPK-A |
| 01272 | p35S-MpCBL-A-LgBiT-t35S<br>pNOS-SmBiT-MpCIPK-B-tNOS<br>pAtUBI10-GUS-tNOS | Split luciferase<br>MpCBL-A-LgBiT<br>SmBiT-MpCIPK-B |
| 01273 | p35S-MpCBL-B-LgBiT-t35S<br>pNOS-SmBiT-MpCIPK-B-tNOS<br>pAtUBI10-GUS-tNOS | Split luciferase<br>MpCBL-B-LgBiT<br>SmBiT-MpCIPK-B |
| 01274 | p35S-MpCBL-C-LgBiT-t35S<br>pNOS-SmBiT-MpCIPK-B-tNOS<br>pAtUBI10-GUS-tNOS | Split luciferase<br>MpCBL-C-LgBiT<br>SmBiT-MpCIPK-B |
| 01275 | p35S-AtCBL4-LgBiT-t35S<br>pNOS-SmBiT-MpCIPK-B-tNOS<br>pAtUBI10-GUS-tNOS | Split luciferase<br>AtCBL4-LgBiT<br>SmBiT-MpCIPK-B |
| 01276 | p35S-MpCBL-A-LgBiT-t35S<br>pNOS-SmBiT-MpCIPK-A-ΔNAF-tNOS<br>pAtUBI10-GUS-tNOS | Split luciferase<br>MpCBL-A-LgBiT<br>SmBiT-MpCIPK-A ΔNAF |
| 01277 | p35S-MpCBL-B-LgBiT-t35S<br>pNOS-SmBiT-MpCIPK-A-ΔNAF-tNOS<br>pAtUBI10-GUS-tNOS | Split luciferase<br>MpCBL-B-LgBiT<br>SmBiT-MpCIPK-A ΔNAF |
| 01278 | p35S-MpCBL-C-LgBiT-t35S<br>pNOS-SmBiT-MpCIPK-A-ΔNAF-tNOS<br>pAtUBI10-GUS-tNOS | Split luciferase<br>MpCBL-C-LgBiT<br>SmBiT-MpCIPK-A ΔNAF |
| 01279 | p35S-AtCBL4-LgBiT-t35S<br>pNOS-SmBiT-MpCIPK-A-ΔNAF-tNOS<br>pAtUBI10-GUS-tNOS | Split luciferase<br>AtCBL4-LgBiT<br>SmBiT-MpCIPK-A ΔNAF |
| 01280 | p35S-MpCBL-A-LgBiT-t35S<br>pNOS-SmBiT-MpCIPK-B-ΔNAF-tNOS<br>pAtUBI10-GUS-tNOS | Split luciferase<br>MpCBL-A-LgBiT<br>SmBiT-MpCIPK-B ΔNAF |
| 01281 | p35S-MpCBL-B-LgBiT-t35S | Split luciferase |

| | | |
|---|---|---|
| | pNOS-SmBiT-MpCIPK-B-ΔNAF-tNOS<br>pAtUBI10-GUS-tNOS | MpCBL-B-LgBiT<br>SmBiT-MpCIPK-B ΔNAF |
| 01282 | p35S-MpCBL-C-LgBiT-t35S<br>pNOS-SmBiT-MpCIPK-B-ΔNAF-tNOS<br>pAtUBI10-GUS-tNOS | Split luciferase<br>MpCBL-C-LgBiT<br>SmBiT-MpCIPK-B ΔNAF |
| 01283 | p35S-AtCBL4-LgBiT-t35S<br>pNOS-SmBiT-MpCIPK-B-ΔNAF-tNOS<br>pAtUBI10-GUS-tNOS | Split luciferase<br>AtCBL4-LgBiT<br>SmBiT-MpCIPK-B ΔNAF |
| 01284 | p35S-MpCBL-A-LgBiT-t35S<br>pNOS-SmBiT-AtCIPK24-tNOS<br>pAtUBI10-GUS-tNOS | Split luciferase<br>MpCBL-A-LgBiT<br>SmBiT-AtCIPK24 |
| 01285 | p35S-MpCBL-B-LgBiT-t35S<br>pNOS-SmBiT-AtCIPK24-tNOS<br>pAtUBI10-GUS-tNOS | Split luciferase<br>MpCBL-B-LgBiT<br>SmBiT-AtCIPK24 |
| 01286 | p35S-MpCBL-C-LgBiT-t35S<br>pNOS-SmBiT-AtCIPK24-tNOS<br>pAtUBI10-GUS-tNOS | Split luciferase<br>MpCBL-C-LgBiT<br>SmBiT-AtCIPK24 |
| 01287 | p35S-AtCBL4-LgBiT-t35S<br>pNOS-SmBiT-AtCIPK24-A-tNOS<br>pAtUBI10-GUS-tNOS | Split luciferase<br>AtCBL4-LgBiT<br>SmBiT-AtCIPK24 |
| 01247 | p35S-HYG-tNOS<br>pMpEF1α-NLS-pcoCas9-t35S<br>pMpU6-1-sgRNA-MpCIPK-B1-tRNAP<br>pMpU6-1-sgRNA-MpCIPK-B2-tRNAP | CRISPR/Cas9<br>MpCIPK-B knockout |
| 01135 | pNOS-KAN-tNOS<br>p35S-MpCBL-A-VYNE-t35S<br>pAtUBI10-dsRED-tNOS | BiFC<br>MpCBL-A-Venus$^{N}$ |
| 01188 | pNOS-KAN-tNOS<br>p35S-MpCBL-B-VYNE-t35S<br>pAtUBI10-dsRED-tNOS | BiFC<br>MpCBL-B-Venus$^{N}$ |
| 01137 | pNOS-KAN-tNOS<br>p35S-MpCBL-C-VYNE-t35S<br>pAtUBI10-dsRED-tNOS | BiFC<br>MpCBL-C-Venus$^{N}$ |
| 01138 | pNOS-KAN-tNOS<br>p35S-VYCE(R)-MpCIPK-A-t35S<br>pAtUBI10-dsRED-tNOS | BiFC<br>Venus$^{C}$-MpCIPK-A |
| 01139 | pNOS-KAN-tNOS<br>p35S-VYCE(R)-MpCIPK-B-t35S<br>pAtUBI10-dsRED-tNOS | BiFC<br>Venus$^{C}$-MpCIPK-B |
