## Supplemental Figure 1 for "CIPK-B is essential for salt stress signalling in *Marchantia polymorpha*"

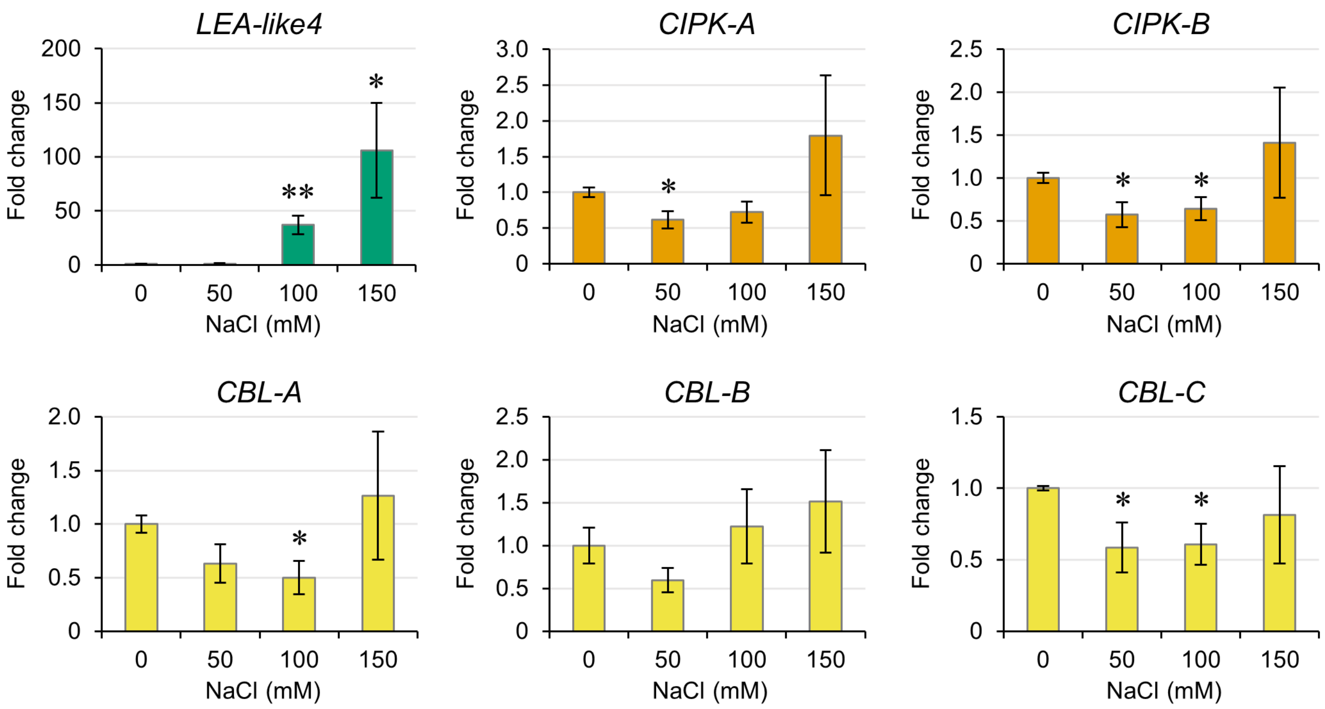


**Figure S1: Regulation of *CBL* and *CIPK* gene expression by salt stress in Tak-1 plants.** Expression changes of the indicated genes were measured by RT-qPCR in thallus tissue of Tak-1 plants grown in the presence of the indicated concentrations of NaCl for 7 days. Data represent mean ± standard error from three independent biological replicates. Significance in a pairwise two-tailed t-test relative to 0 mM treatment is indicated at $p<0.05$ (one star) or $p<0.01$ (two stars).
