## Supplemental Figure 2 for "CIPK-B is essential for salt stress signalling in *Marchantia polymorpha*"

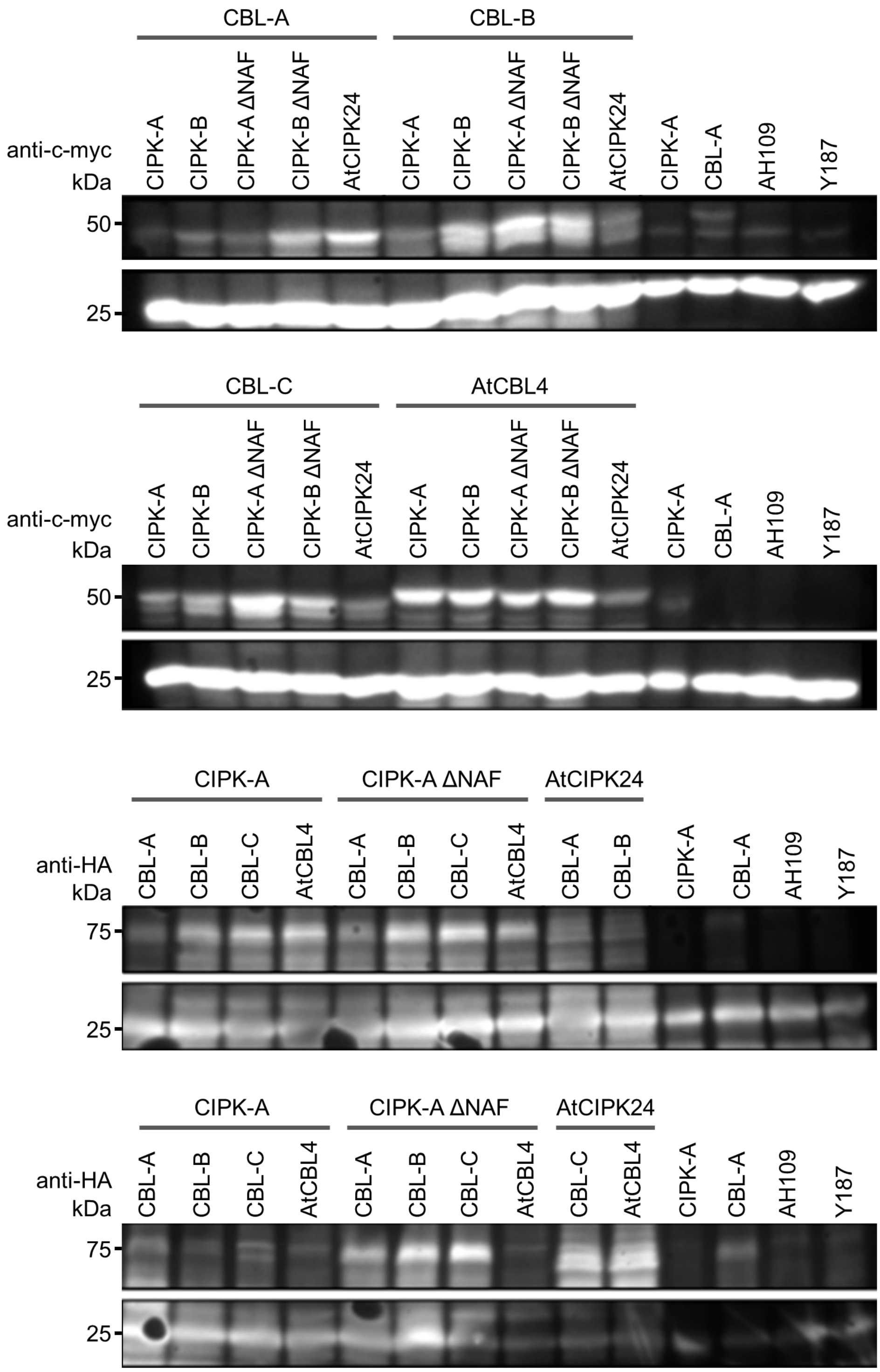


**Figure S2: Western blotting confirms expression of all *M. polymorpha* CBLs and CIPKs in strains used for yeast two-hybrid assays.** Western blotting using c-myc and HA antibodies identified proteins of the expected size for GAL4-BD-3xc-myc-CBLs and GAL4-AD-3xHA-CIPKs, respectively (upper panels). Expected sizes of the GAL4-BD-3xc-myc-CBLs are: 48.5 (MpCBL-A), 46.6 (MpCBL-B and MpCBL-C) and 47.8 kDa (AtCBL4). Expected sizes of the GAL4-AD-3xHA-CIPKs are: 69.7 (MpCIPK-A), 70.0 (MpCIPK-B), 67.1 (MpCIPK-A ΔNAF), 67.4 (MpCIPK-B ΔNAF) and 70.2 kDa (AtCIPK24). Lower panels correspond to loading control, using native yeast peroxidase (PRX1) activity as a proxy for total protein (expected size of 25 kDa).
