## Supplemental Figure 3 for "CIPK-B is essential for salt stress signalling in *Marchantia polymorpha*"

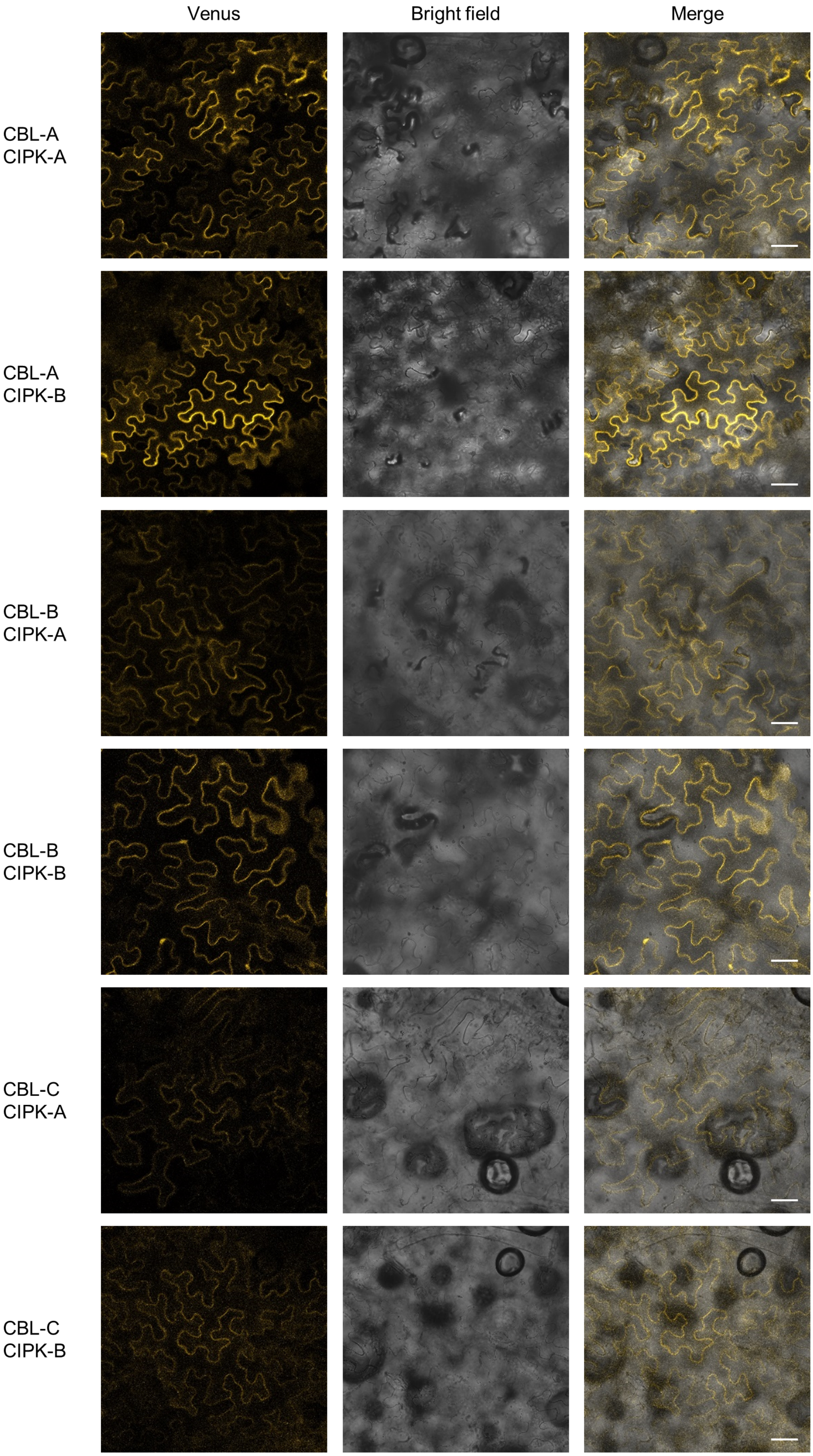


**Figure S3: Bimolecular fluorescence complementation confirms all *M. polymorpha* CBLs and CIPKS interact in *N. benthamiana*.** Constructs containing the C-terminal half of the Venus fluorescent protein fused to each CIPK (VenusC-CIPK) and the N-terminal half of Venus fused to each CBL (CBL-VenusN) were transiently expressed in *N. benthamiana* leaf epidermal cells. Venus fluorescence, bright field and merged images are shown for each *M. polymorpha* CBL-CIPK interaction. Scale bar in all panels = 100 µm.
